# A robust and fast two-sample test of equal correlations with an application to differential co-expression

**DOI:** 10.1101/2022.01.23.477409

**Authors:** Liang He, Ian Philipp, Stephanie Webster, Alexander M. Kulminski

## Abstract

A robust and fast two-sample test for equal Pearson correlation coefficients (PCCs) is important in solving many biological problems, including, e.g., analysis of differential co-expression. However, few existing methods for this test can achieve robustness against deviation from normal distributions, accuracy under small sample sizes, and computational efficiency simultaneously. Here, we propose such a method for testing DIfferential COrrelation using a Saddlepoint Approximation of the Residual bootstrap (DICOSAR). To achieve robustness, accuracy, and efficiency, DICOSAR combines the ideas underlying the pooled residual bootstrap, the signed root of a likelihood ratio statistic, and a multivariate saddlepoint approximation. Through a comprehensive simulation study and a real data analysis of gene co-expression, we demonstrate that DICOSAR is accurate and robust in controlling the type I error rate for detecting differential correlation and provides a faster alternative to the permutation method. We further show that it can also be used for testing differential correlation matrices. These results suggest that DICOSAR provides an analytical approach to facilitate rapid testing for the equality of PCCs in a large-scale analysis.

## Introduction

Testing the equality of Pearson correlation coefficients (PCCs) between two groups is one of the most fundamental statistical problems for investigating whether the dependency between variables differs between groups of interest. Its application can be widely found in many research areas, including biology and social science. For example, the correlation between gene expression can imply co-regulation in the same pathway and thus provide insights into the study of dysfunctional regulatory networks (de la Fuente, 2010).

Despite its importance, to the best of our knowledge, this two-sample homogeneity test of PCCs still poses significant challenges if pursue robustness, statistical accuracy, and computational efficiency are simultaneously required. The major challenges in real data analysis include small sample sizes, violation of a normality assumption, and computational burden. The computational cost often becomes a major concern in applications involving a considerable number of tests. For example, in the co-expression analysis, millions of gene pairs may need to be tested. A common fast approach for testing PCCs is through Fisher’s z-transformation (Fisher, 1925), which converts the sample distribution of the PCC to a normal distribution with the variance equal to 1/(*n* – 3), where *n* is the sample size. The two-sample homogeneity test can then be readily carried out by testing a difference between two variables of normal distributions. Unfortunately, the normality of the z-transformation is valid only under the strong assumption that the variables are bivariate normal (Hawkins, 1989), which can be easily violated in real data analysis. Multiple studies demonstrate that the distribution of the z-transformation departs from a normal distribution if such an assumption does not hold (Bishara and Hittner, 2017; Puth et al., 2014). Another widely used approach is to first transfer the original variables before testing the correlation. However, as shown in (Bishara and Hittner, 2017), many transformations of the raw data that aim to approach normality might not completely solve the problem or cannot be used if the linear relationship must be measured on the original scale. On the other hand, rank-based transformations like Spearman’s rho can reduce the statistical power if the normality does hold for the raw data (Pernet et al., 2013).

To relax the normality assumption, Hawkins (Hawkins, 1989) proposes a delta method based on U-statistics to obtain the asymptotic distribution of the z-transformation and shows that the variance depends on higher-order joint moments of the two variables. The problem of this method is that the sample higher-order joint moments are less accurate under a small sample size. Nevertheless, as shown in our simulation study, the delta method exhibits an inflated type I error rate under a small or even moderate sample size of 200 subjects, which is even worse than the z-transformation for bivariate normal variables. In, e.g., gene expression data, it is very common to have only dozens of samples in a group, and therefore such inflation is not ignorable in many real data analyses. Instead of directly estimating the joint moments, an approximation distribution is developed and shows better accuracy in terms of confidence interval (Bishara et al., 2018). However, our simulation indicates that its performance depends on the PCC and the underlying distribution of the variables. Consequently, various resampling strategies such as the residual permutation or bootstrap (Boos and Brownie, 1989; Krzanowski, 1993; Tesson et al., 2010; Yang and DeGruttola, 2012; Zhang and Boos, 1992, 1993) are broadly adopted to compute the p-value in real problems. Despite being robust against assumptions and its straightforward implementation, these resampling methods can be computationally expensive particularly in the multiple testing problem. To obtain accurate significant p-values, the resampling methods need to generate a huge number of replicates, which is computationally inhibitive, particularly in a situation where many hypotheses need to be tested. For example, >10^6^ random samples might be required for providing a decent estimate of a p-value <10^−5^. Therefore, it is appealing to find a fast, accurate, and robust method for testing the equality of two PCCs that can work well even under a small sample size and does not rely on a resampling procedure.

The aim of this study is to develop such an analytical algorithm for testing the equality of PCCs that can accurately control type I errors even under a small sample size and nonnormal distributions. We propose a method for testing DIfferential COrrelation using a Saddlepoint Approximation of the Residual bootstrap, referred to as DICOSAR. DICOSAR combines the ideas underlying the residual bootstrap method (Zhang and Boos, 1992, 1993) and an accurate approximation for the cumulative distribution of a function of multiple random variables proposed in (DiCiccio et al., 1994). Our basic idea is using a multivariate saddlepoint method (Daniels and Young, 1991) to approximate the distribution of the summary statistics under the null hypothesis and then employing a higher-order approximation for the cumulative distribution of a smooth function of the summary statistics (DiCiccio and Martin, 1991). Under the same sample size, we show that this method is much more accurate than the delta method, which assumes normality in both steps. In a comprehensive simulation study and an analysis of differential gene co-expression, we demonstrate that DICOSAR has comparable performance to the pooled residual permutation in controlling the type I errors, and is computationally faster than the permutation method.. To demonstrate its performance in real data analysis, we applied DICOSAR to detect genes showing differential co-expression with *APOE* between controls and patients with Alzheimer’s disease (AD) in bulk and single-nucleus RNA-seq (snRNA-seq) data sets.

## Materials and Methods

### The main algorithm in DICOSAR

We start with reviewing the residual permutation and bootstrap strategies. Consider that we collect quantitative data in two groups for which we want to test the equality of the PCCs of two continuous variables of interest. Denote the datasets of the two groups by 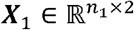 and 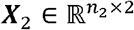, where *n*_1_ and *n*_2_ are the sample size, respectively. By centering and standardizing the variables within each group, we obtain the residuals 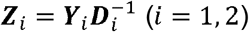, where 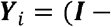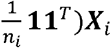 and 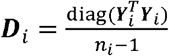. Here, ***I*** is the identity matrix, **1** is the *n*_*i*_ × 1 matrix of ones, the subscript *T* stands for the matrix transpose, and diag(**M**) is a diagonal matrix containing the diagonal entries of **M**. The statistic that we propose to test the equality of the PCCs is

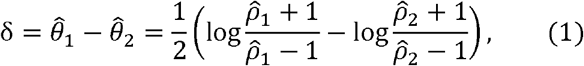

where 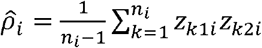 is the sample PCC *ρ*_*i*_ in group *i* and *z*_*kji*_ is the element in ***Z***_*i*_ corresponding to the *j*^th^ variable of sample *k* in group *i*. Throughout this manuscript, matrices or vectors are denoted by boldface uppercase letters. The statistic δ is essentially the difference of the Fisher’s z-transformation between the two groups. To test *ρ*_1_ = *ρ*_2_, we need the sampling distribution of the statistic δ under the null hypothesis. Note that 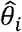 follows a normal distribution only when ***X***_*i*_ follow a bivariate normal distribution. A robust approach to obtain the null distribution without the strong assumption of normality is the pooled residual permutation (Krzanowski, 1993; Tesson et al., 2010) or bootstrap (Yang and DeGruttola, 2012; Zhang and (Krzanowski, 1993; Tesson et al., 2010) or bootstrap (Yang and DeGruttola, 2012; Zhang and with or without replacement. The rationale of such a pooling procedure is that under the null with or without replacement. The rationale of such a pooling procedure is that under the null fourth moment of the sample distribution (Zhang and Boos, 1993). So, one can generate a pooled sample ***Z*** by stacking the rows of ***Z***_1_ and ***Z***_2_ and then resampling from the rows of ***Z***. In each random sample ***Z***\* the dataset is split into 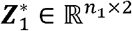 and 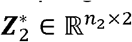, and δ* is calculated according to formula (1) by substituting the original data with 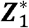 and 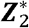. Our simulation study shows that this strategy is robust against deviation from the normality assumption and controls type I errors properly. More consideration about violation of the fourth-moment condition can be found in the Discussion section. However, the drawback of the permutation or bootstrap method is its computational intensity, particularly for testing many pairs of variables. In this case, it may require a very large number of permutations to obtain a significant p-value that passes the multiple testing correction.

The key idea in DICOSAR is to obtain an accurate analytical approximation of the cumulative null distribution of δ without resorting to a time-consuming resampling procedure. Following the spirit of the pooled residual bootstrap method, the distribution of the statistic δ under the null hypothesis can be obtained based on the pooled residual sample ***Z***. That is, ***Z*** is treated as a sample from the null hypothesis. More specifically, *n*_1_ and *n*_2_ samples are randomly chosen from ***Z*** independently for the two groups, denoted by 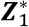 and 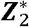. Then we have

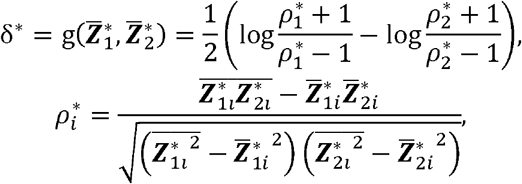

where 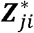, is the *j*^th^ variable (*j* ϵ {1,2}) in group *i* and 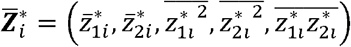 is a vector of the summary statistics including the sample means 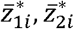, second moments 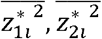, and joint moment 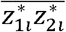 for group *i*. Thus, it remains to derive the distribution of δ* based on the joint distribution of the summary statistics 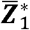 and 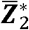.

Following the spirit of the analytical approximation to bootstrap distribution functions proposed in (DiCiccio et al., 1994), we approximate the distribution of δ* in two steps. In the first step, we approximate the joint distribution of 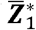 and 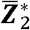 using a multivariate saddlepoint method. Because the two groups are independent, we can apply the saddlepoint method to 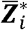 separately. More specifically, the cumulant generating function (CGF) of the joint distribution of 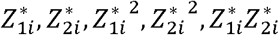 conditional on ***Z***, which is independent of *i*, is

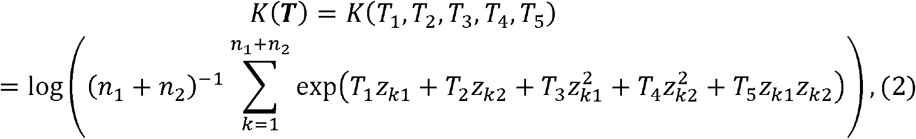

where *z*_*kj*_ is the element at the *k*^th^ row and *j*^th^ column in ***Z***. Then, the general multivariate saddlepoint approximation (Butler, 2007; Daniels and Young, 1991) to the joint distribution of 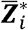 given ***Z*** is

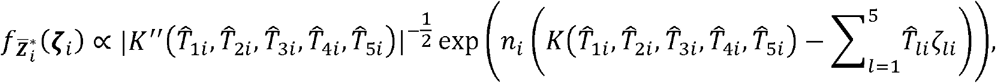

where 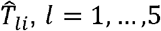, satisfy the following saddlepoint equation

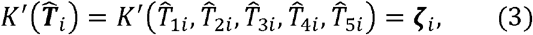

where *K*′ and *K*′′ are the Jacobian and Hessian matrix of the CGF, respectively. Under the assumption that the two groups are independent, the joint distribution of 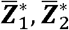 is approximated by

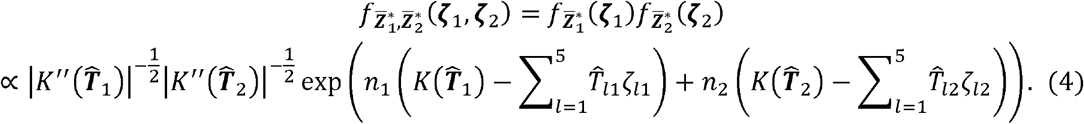

Our goal is to approximate the tail probability 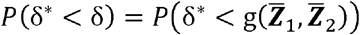, Given the approximated joint distribution 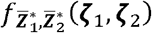, in the second step, we attempt to approximate *P*(δ* < δ) using a signed root of the likelihood ratio statistic, which has been discussed in (Barndorff-Nielsen, 1986; McCullagh, 1984). Let

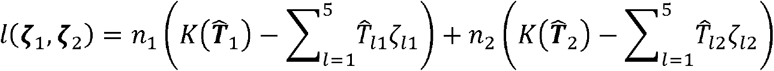

and

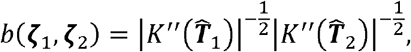

where 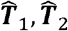 are functions of **ζ**_1_, **ζ**_2_ through the saddlepoint equation (3). We define the signed We define the signed statistic as

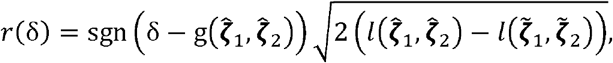

where sgn(·) is the sign function extracting the sign of a real number, 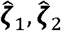 are the values that maximize *l*(**ζ**_1_, **ζ**_2_), and 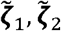 are the values that maximize *l*(**ζ**_1_, **ζ**_2_) subject to the constraint g(**ζ**_1_, **ζ**_2_) = δ. To find 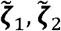 under this nonlinear constraint, we introduce the following Lagrangian

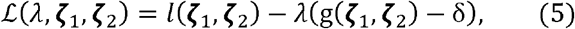

where *λ* is the Lagrange multiplier. Thus, 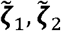 can be obtained by solving the equations of the gradient of the Lagrangian, i.e., 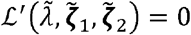. Denote 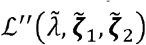 the bordered Hessian matrix of the Lagrangian evaluated at 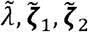. We adopt the following high-order tail probability approximation proposed in (DiCiccio and Martin, 1991),

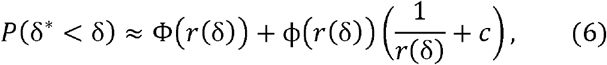

where Φ and ϕ are the cumulative and density distribution functions of the standard normal distribution, respectively,

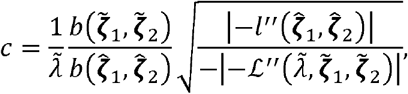

and 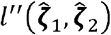 is the Hessian matrix of *l*(**ζ**_1_, **ζ**_2_) evaluated at 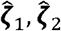. This approximation (6) is derived in (DiCiccio and Martin, 1991) by combining the two approximation methods proposed in (Diciccio et al., 1990) and (Tierney et al., 1989, 1991). In the expression of *c*, 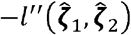 is positive definite at the minimum 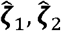 and the determinant of the minus bordered Hessian matrix 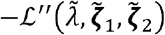 is negative if 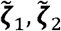 is a maximum because the sign of 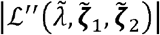 is 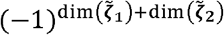 based on the rule of the second derivative test for constrained local extrema (see e.g., (Colley, 2006)). Therefore, the term in the square root is always positive if the constrained optimization algorithm finds the correct solution. Finally, by substituting the approximation (6), the p-value for a two-sided test of *ρ*_1_ = *ρ*_2_ can be obtained by

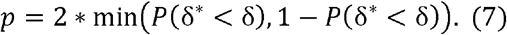

### Computational implementation and numerical issues

The major computational burden in DICOSAR is to solve the saddlepoint equations in (3) to obtain 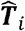 and the Lagrangian equations 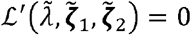 to obtain 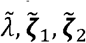. We use the *multiroot* function in the rootSolve R package to solve the equations in (3) numerically. Because 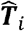 maximize *l*(**ζ**_1_, **ζ**_2_) given *ζ*_*i*_ to solve the 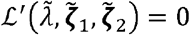, it follows from the envelope theorem (see e.g., (Carter, 2001)) that

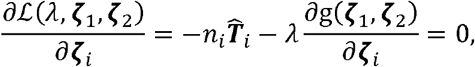

and

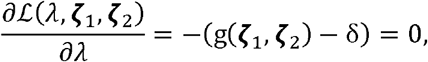

where 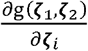 is the partial derivative with respect to *ζ*_*i*_ and can be calculated explicitly. We use the nleqslv function with the parameters “method=;‘Newton’” and “global=‘hook’” to solve these 11 equations numerically. We use the jacobian function in the numDeriv R package for computing Jacobian matrices numerically. Practically, we find that the algorithm converges very well except for some rare cases where the matrix 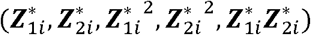 is almost singular. The higher-order approximation (6) is very accurate in general, but *c* might be sensitive to the numerical precision of the *jacobian* function when 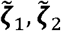 are very close to 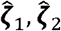, In this situation, 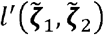 is almost zero and thus 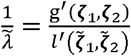 becomes less accurate and stable, Therefore, when 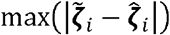 is very small (e.g., <0.001), we practically adopt the following first-order approximation, which is 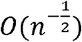,

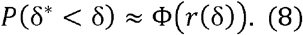

Additionally, special attention should be paid when applying this method to discrete random variables, especially if they have only several levels, e.g., genotypes. For example, if one of the variables has only two values, zero and one, the conditional distribution of 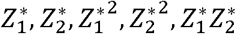 given ***Z*** is degenerated because 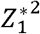 (or 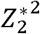) is determined by 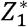 (or 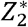). The similar issue occurs if the rows of ***Z*** have only five or less different levels. In these situations, this method cannot be applied directly without a specific adjustment for the data.

### Comparison with other methods

We consider three analytical and resampling methods, and compare their statistical and computational performance with DICOSAR. First, we include the pooled residual permutation method. In this method, we merge the standardized residuals to generate ***Z*** and permute the rows of ***Z*** for *M* times. In our simulation study, we chose *M* to be 5000, and in our real data analysis, we ran *M* times until there were at least five more extreme values than the observed PCC. In each of the permutation replicates, we split the permuted data into two groups and calculate the statistic in formula (1). We obtain an empirical null distribution from the *M* replicates and calculate the p-value using equation (7).

We also consider a fast testing algorithm based on the Delta method proposed in (Hawkins, 1989). In this method, we assume that 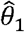 and 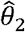 in the z-transformation (1) follow normal distributions with means 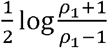 and 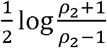, and variances 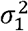 and 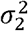 in these two groups. Hence, under the null hypothesis of *ρ*_1_ = *ρ*_2_, the difference 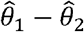 follows a zero-mean normal distribution with variance 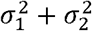. An asymptotic estimate of 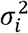 using the Delta method is a function of the fourth moment if *Z*_1*i*_ and *Z*_2*i*_, and the joint moments 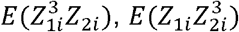, and 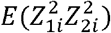, where *Z*_*ji*_ is the *j*^th^ variable in group *i*. We use the sample moments to estimate these quantities. Because the sample joint moments might not be accurate estimates under a small sample size, (Bishara et al., 2018) propose an improved method by assuming third-order polynomials for the variables to estimate these joint moments, which shows superior performance in terms of estimating confidence intervals. We further include a variant of the Delta method introduced in (Bishara et al., 2018). To run this method, we directly use the R script provided in the supplemental material of (Bishara et al., 2018).

We further include a separate bootstrap method, in which the variances of 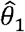 and 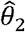 are estimated using a non-parametric bootstrap within each of the groups. In the simulation study, we tested 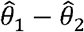 by assuming that it follows a zero-mean normal distribution under the null hypothesis using 2000 bootstrap replicates.

### Global test of multiple differential correlation coefficients

In some applications, one can be further interested in testing the global correlation pattern of multiple (>2) variables after testing each pair of these variables. For example, given *K* variables, one may want to test the equality of two correlation matrices ***R***_1_ and ***R***_2_ ϵ ℝ^*K*×*K*^, or a subset of the elements in the correlation matrices. Suppose that we perform a global test of all *K*(*K* – 1)/2 elements in the *K* × *K* correlation matrices. One simple analytical approach is to combine the *K*(*K* – 1)/2 p-values obtained by testing each pair of the *K* variables. Because these p-values are not independent, we adopt the Cauchy combination test (Liu and Xie, 2020). The idea underlying this test is based on the finding that a weighted sum of some class of correlated Cauchy variables still follows a Cauchy distribution as proved by (Pillai and Meng, 2016). Specifically, let *p*_*kl*_ be the p-value of testing the equality of PCCs between the *k*th and *l*th variables. The test statistic is

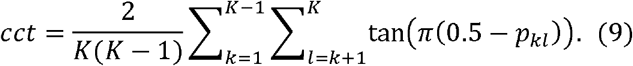

Under the null hypothesis of the equality of the two correlation matrices, *cct* approximately follows a standard Cauchy distribution at the extreme tail under a large sample size. The p-value for testing small *p*_*kl*_ globally is calculated by the cumulative probability of a standard Cauchy variable from the right tail, i.e.,

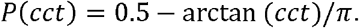

This test works well for aggregating a small number of p-values. If the dimension is large, additional assumptions about the correlation structure of these single tests are required (Liu and Xie, 2020). Roughly speaking, the single tests cannot be too closely correlated with each other at a large scale under the high-dimensional scenario.

### Simulation study

We perform a comprehensive simulation study to investigate the statistical and computational performance of DICOSAR and compare it with the other methods. We evaluate the empirical type I error rate under a wide range of settings that differ in the sample size, the distribution of the variables, and the correlation strength. Specifically, the *j*^th^ simulated data set of group *i* is produced using the following generative model

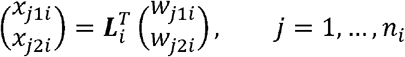

where ***L***, is the Cholesky decomposition of the correlation matrix, i.e.,

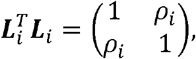

and *w*_*j1i*_ and *w*_*j2i*_ are independent and identically distributed random variables. Here, we evaluate the following distributions for *w*_*j1i*_ and *w*_*j2i*_, (i) the standard normal distribution, (ii) two t-distributions with six and four degrees of freedom, respectively, (iii) the gamma distribution with the shape and rate equal to 1, and (iv) a mixture of a standard normal distribution and a normal distribution with mean equal to five. The rationale of choosing these distributions for the assessment is to examine different scenarios, including skewness, excessive kurtosis, and multimodal distributions. The two t-distributions are heavy-tailed distributions, one with a finite fourth moment and the other with an infinite fourth moment. The gamma distribution has both skewness and excessive kurtosis, and the mixture of two normal distributions is a common bimodal distribution. We consider *n*_*i*_, the sample size of group *i*, being 25, 50, 100, 200, and 400, and *ρ*_*i*_ being 0, 0.4, and 0.8 for independent, moderate, and strong correlations, respectively. We investigated the empirical power under the same settings except that only the normal and gamma distributions are considered.

We examine the empirical type I error rate of testing the equality of two correlation matrices by using statistic (9). Similarly, we simulate the *K*-dimensional data set of group *i* using the generative model

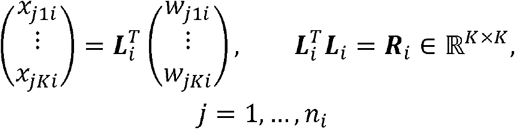

where ***R***_*i*_, is the correlation matrix of group *i*. Here, we consider three correlation patterns for ***R***_*i*_, including an identity matrix, an autoregressive correlation matrix

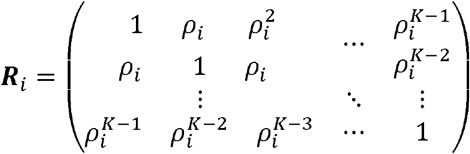

with *ρ*_*i*_ = 0.5, and a matrix whose off-diagonal elements share the same value, i.e., 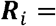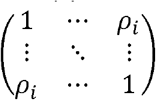 with *ρ*_*i*_ being 0.3 and 0.6.

### Processing of gene expression data

In the real data analysis, we applied DICOSAR to co-expression analyses using two gene expression data sets in ROSMAP. In both analyses, we used the diagnosis of AD based on brain pathology to define the control and AD groups.

The raw count matrix of the bulk RNA-seq gene expression data of 482 samples in ROSMAP (Bennett et al., 2012a, 2012b) was downloaded from Synapse. The biological samples are extracted from the human dorsolateral frontal cortex. Gene-level quantification is conducted by RSEM (Li and Dewey, 2011). More details about the sample information and data generation can be found in (Bennett et al., 2012a, 2012b). For each sample, we normalized the raw counts by dividing by the total library size (i.e., summing up the count of each gene) of the sample followed by taking the logarithm transformation. To avoid zeros for the logarithm, we added 0.5 as the pseudo-count before normalizing the counts. This transformed data set was used for testing the differential co-expression with *APOE*.

We downloaded the 48-sample snRNA-seq raw count data set (Mathys et al., 2019) in ROSMAP from synapse. After the quality control, this data set contains 70,634 cells and 17,926 genes in the human frontal cortex. For each of the neural cell types, we generated a pseudo-bulk count matrix by aggregating all cells of that cell type belonging to the same sample. We adopted the cell-type annotation provided in (Mathys et al., 2019). This leads to a count matrix containing 17,926 genes and 48 samples for each cell type. We normalized the aggregated counts by dividing by the total library size of each sample, which were then used for the differential co-expression analysis.

### Data and code availability

This manuscript was prepared using limited access datasets obtained through Synapse (https://www.synapse.org/#!Synapse:syn3219045; https://www.synapse.org/#!Synapse:syn18485175). The program used to analyze the simulated data and the gene expression data can be found in the dicosar R package at https://github.com/lhe17/dicosar.

## Results

### DICOSAR has similar performance to permutation in controlling type I errors

We evaluated the performance of DICOSAR in controlling type I error rate under various settings. For comparison, we included the pooled residual permutation, the Delta method, the improved Delta method and the bootstrap method, which are described in detail in the Methods section. We also included a simplified version of DICOSAR using the approximation (8) instead of (6) to assess how much improvement can be achieved by using the higher-order approximation for the distribution of the signed root of the likelihood ratio statistic. In the scenario where the two variables are generated from a linear transformation of independent normal variables (i.e., a bivariate normal distribution), all methods control the type I errors well at the significance level of 5% under the largest sample size (i.e., 400 subjects per group) (Fig. 1A), which is not surprising because the assumption of normality holds under a large sample size. The correlation strength between the two variables seems to have little impact on the empirical type I error rate. Whereas both DICOSAR and the permutation control the type I errors under all these settings, we start to see noticeable inflation of the type I errors from the Delta method when the sample size per group drops to 100. The empirical type I error rate of the Delta method is above 10% under 25 samples per group. The improved Delta method performs much better than the Delta method under this setting and is comparable to the bootstrap method, but still shows inflation of the type I errors to some extent when the sample size is 25. By comparing DICOSAR and its simplified version, we find that the higher-order approximation for the likelihood ratio statistic (8) has a strong improvement and enables DICOSAR to have almost identical performance of the pooled residual permutation.

**Figure 1:**
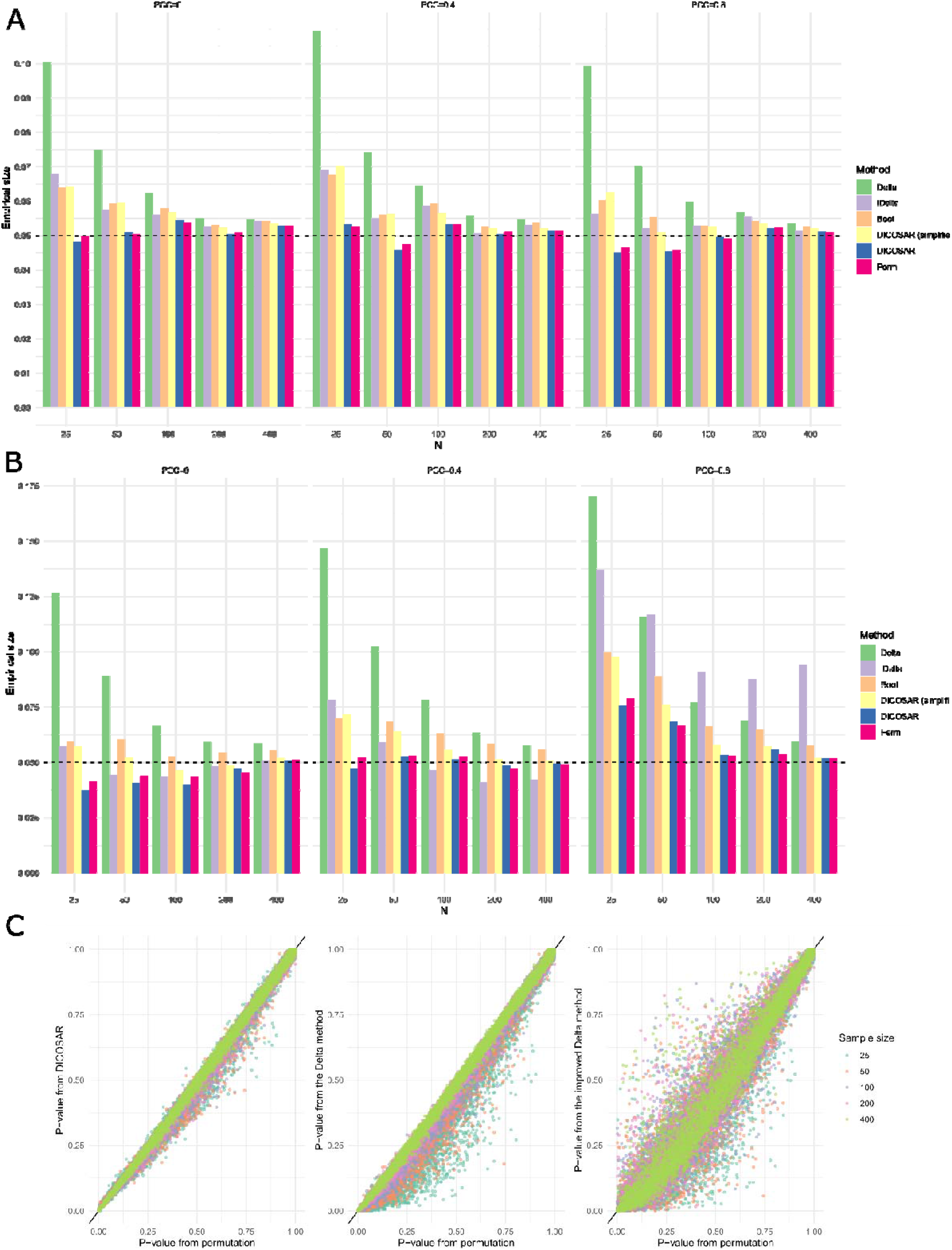
Performance of DICOSAR in controlling the false positive rate for testing the equality of PCCs between two groups. (A) & (B) Empirical type I error rate of six methods at the significance level of 5%. The data are generated from (A) a bivariate normal distribution (B) a linear transformation of independent gamma-distributed variables. Delta: the Delta method; iDelta: the improved Delta method; Boot: the bootstrap method; Perm: the pooled residual permutation; DICOSAR (simplified): a simplified version of DICOSAR using the approximation (8) instead of (6). N: sample size of each group. (C) Comparison of the p-values estimated by the pooled residual permutation with those estimated by DICOSAR, the Delta method, and the improved Delta method.

Next, we investigated the performance when the two variables are generated from a linear transformation of independent variables from non-normal distributions. Variables from non-normal distributions are ubiquitous in real data analysis of e.g., gene expression. Here, we considered heavy-tailed distributions, bimodal distributions, and skewed distributions. We observe a similar overall pattern except for the improved Delta method when the variables are generated from a heavy-tailed t-distribution with six degrees of freedom (d.f.) (Fig. S1A). The improved Delta method fails to control the type I errors when the PCC is large. Among the other methods, we observe slightly higher inflation of type I errors across most scenarios for the Delta method and the bootstrap method than that in the normal case. We then considered a bimodal distribution generated from a mixture of two normal distributions (See the Methods section for more detail). Again, the improved Delta method fails to achieve the expected type I error rate in the cases with a moderate to high correlation, even when the sample size is large (Fig. S1B).

Most of these methods show deflation of the type I errors under the small sample sizes when the PCC is very high. Interestingly, the bootstrap method works very well for this mixture distribution and shows almost the same performance as DICOSAR and the permutation method. The situation becomes slightly different when we consider a gamma distribution with the shape and rate parameters equal to one, which has both skewness and excessive kurtosis. The worst case occurs to very high correlation strength (PCC=0.8), in which neither DICOSAR nor the permutation method can control the type I error rate accurately under a small sample size (<50) although they still work properly under the larger sample sizes (Fig. 1B). The other methods show substantially inflated type I error rate even under the largest sample size.

The consistency of the p-values between DICOSAR and the permutation method is clearly shown in Fig. 1C, where the 5000 p-values in each sample size and method under the gamma distribution with PCC=0.8 are plotted. The points are tightly aligned along the diagonal line, and more consistency is observed under larger sample sizes. In contrast, the Delta method has a clear bias towards smaller values than those from the permutation method, particularly for those small p-values and under small sample sizes (Fig. 1C). Surprisingly, although the improved Delta method has no evident directional bias, the accuracy is very low. For example, as shown in (Fig. 1C), the difference between the improved Delta method and the permutation can be >0.3 for many p-values. These results suggest that DICOSAR shares almost the same performance as the permutation method in all these scenarios, including a sample size as small as 25 per group, demonstrating the remarkable accuracy by using the multivariate saddlepoint approximation and the higher-order approximation for the signed root of the likelihood ratio statistic and thus the robustness against the violation of the normality assumption. On the other hand, the control of type I error rate becomes tough if the data have skewness, heavy tails, a high PCC and small sample size concurrently.

We further examined the accuracy of DICOSAR for approximating the null distribution in the extreme tails (i.e., more significant p-values). The accuracy in the extreme tails is of primary interest in large-scale applications because resampling methods are computationally intensive for estimating significant p-values. Specifically, we investigated the performance of controlling the type I errors at the significance level of 0.1%. The results in Fig. S2 show that DICOSAR controls the type I error rate below its theoretical value of 0.1%. DICOSAR is more conservative than the permutation, particularly when the sample size is small. However, DICOSAR achieves better control of the type I error rate than the permutation under the setting of PCC=0.8, in which the permutation shows substantially inflated type I errors under the gamma distribution and the t-distribution.

### Evaluation of empirical statistical power for detecting differential correlation

Given the robust performance of DICOSAR in controlling the type I errors, we then assess the empirical statistical power for detecting differential correlations. We considered various settings including sample sizes ranging from 25 to 400 samples per group, and different PCCs (small, medium, large). We first investigated a situation in which both variables were generated from bivariate normal distributions. We observe that the empirical power heavily depends on the PCCs in the two groups. If the two PCCs are large, 100 samples per group can achieve ~80% power for detecting a difference of 0.2 between the correlations (PCC=0.8 vs. 0.6) (Fig. 2A). In contrast, when the PCCs in both groups are small or moderate, ~400 samples per group are needed to achieve >80% power for detecting such a difference (Fig. 2A). In the context of gene co-expression analysis, this observation suggests that much fewer samples are needed to detect differential correlations for highly co-expressed genes than uncorrelated genes, which is desirable because highly co-expressed genes in a network or pathway are often of major interest. Nevertheless, for detecting a very small difference (0.05) of correlations, it still requires a very large number of samples (>400) even in the case of high PCCs. We then examined a situation in which the two variables were from a linear transformation of independent gamma-distributed variables. The empirical power in this situation is comparable to that in the case of normal distribution when the PCCs in the two groups are small or moderate. However, the power is substantially reduced when the PCCs are large, compared to the bivariate normal distributions (Fig. 2B).

**Figure 2.**
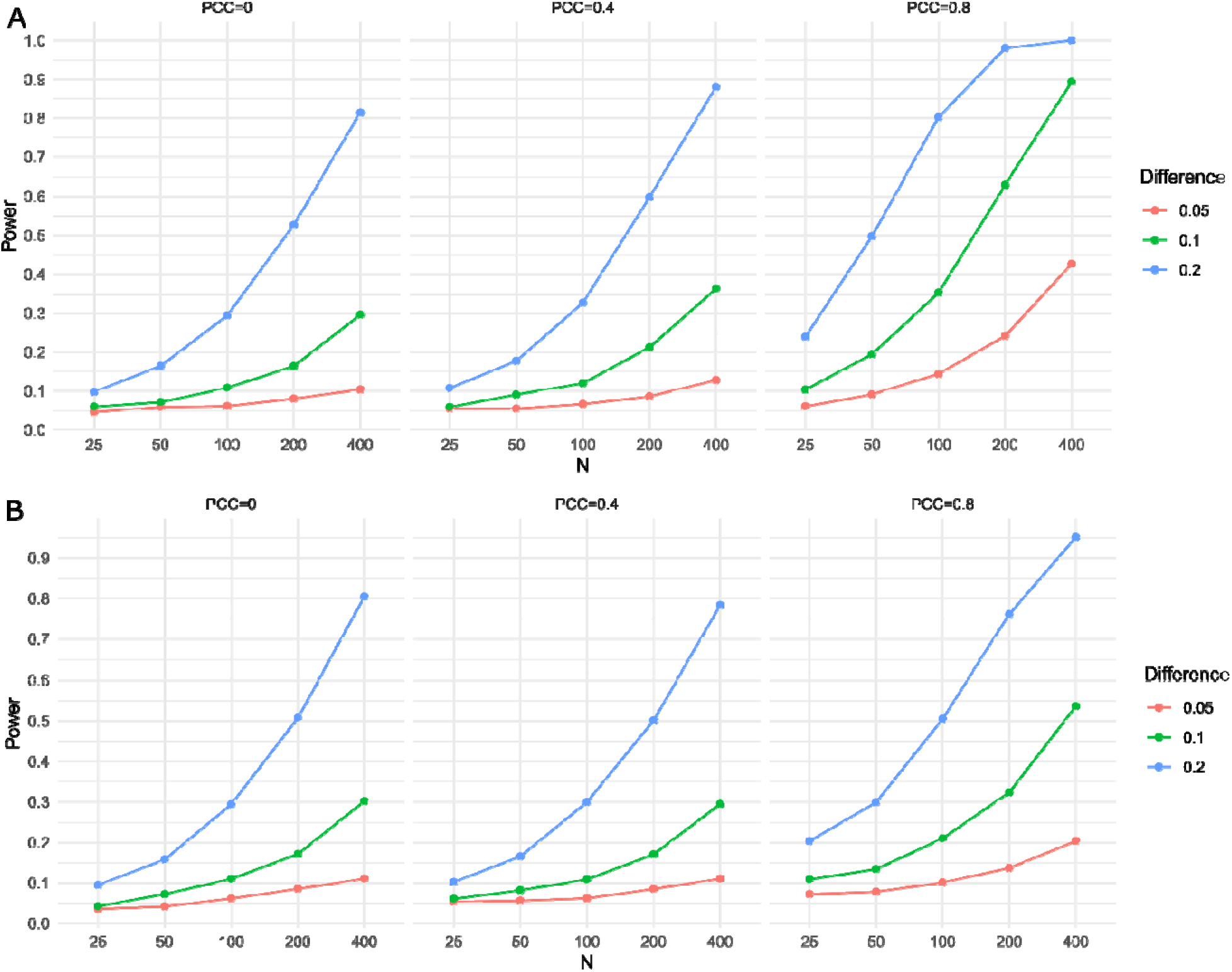
Empirical statistical power of DICOSAR for detecting differential correlation. The data are generated from (A) a bivariate normal distribution (B) a linear transformation of independent gamma-distributed variables. N: sample size of each group. Difference: the difference of PCCs between the two groups. One group has the PCC shown on the top of the panel, and the other has this PCC minus the difference.

### Global test for multiple differential correlations

We next examined the empirical type I error rate of the global test for multiple differential correlations. We focused on an application to test the equality of two *K* × *K* correlation matrices by combining the evidence from all *K*(*K* – 1)/2 single elements in the correlation matrices using statistic (9). We considered various sample sizes per group, ranging from 25 to 200, and *K* = 10 and 50 to cover both low-dimensional and high-dimensional data. In each of these scenarios, we evaluated three correlation patterns, including (i) a mutual independence structure, (ii) an autoregressive correlation structure in which the correlation decays exponentially with the lag, (iii) a correlation matrix in which every pair of two variables share the same PCC. In the third pattern, we considered the PCC being 0.3 and 0.6 for moderate and high correlations, respectively. The simulated data were generated from a multivariate normal distribution. The empirical type I error rate was evaluated at the significance level of 5% and estimated from 1000 random replicates. Fig. 3 shows that the type I error rate was controlled below its theoretical threshold in most scenarios. We observe a deflation of type I errors when the sample size is small (≤50). The higher dimension of the correlation matrices also led to more conservative p-values. We observe that statistic (9) deviates the standard Cauchy distribution when the correlations in the matrices are strong (Fig. S3) although the tail probability is little affected (Fig. 3). Overall, these results indicate that the global test can control the type I error rate but might suffer from some power loss under a small sample size.

**Figure 3.**
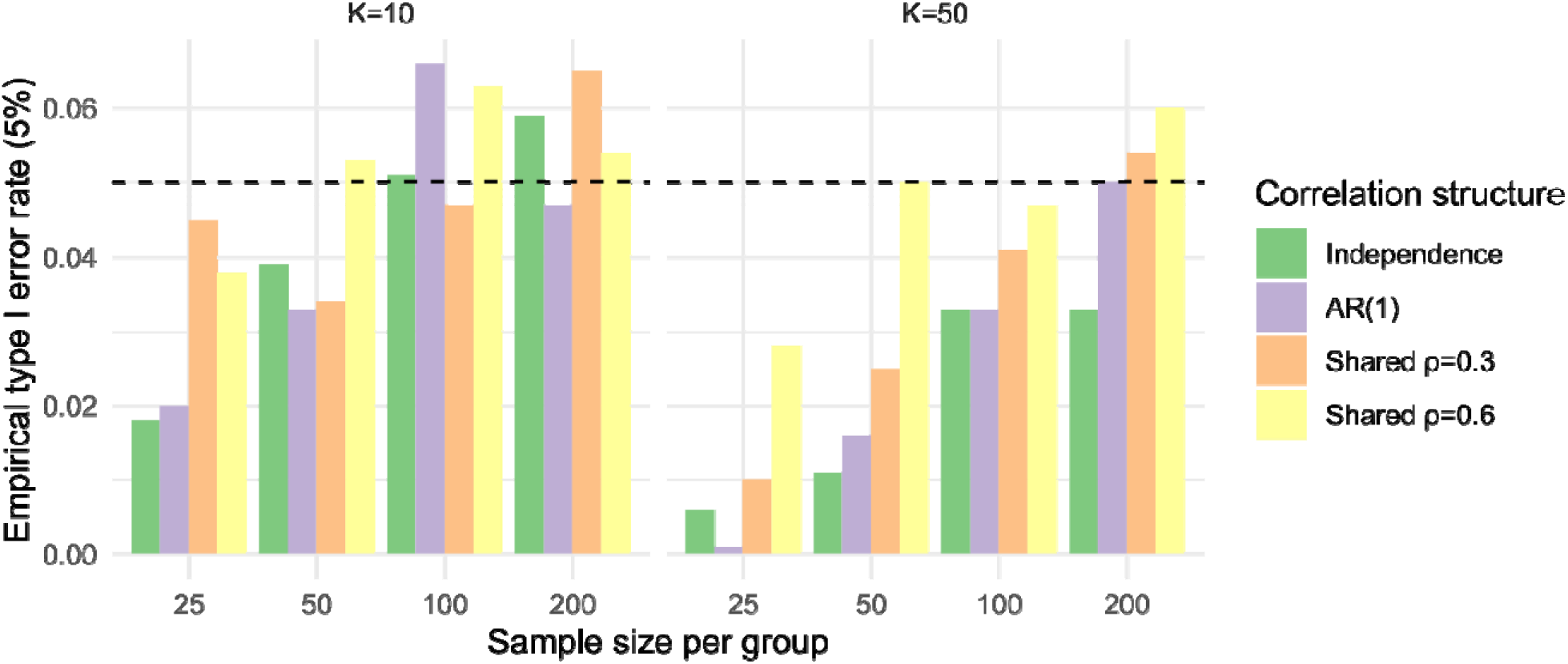
Performance of DICOSAR in controlling the type I error rate for testing the equality of two correlation matrices. Empirical type I error rate is evaluated at the significance level of 5%. The data are generated from a multivariate normal distribution. Dimension (K): the dimension of the correlation matrices. ρ: the value of the off-diagonal elements of the correlation matrices.

### Detecting differential co-expression using DICOSAR

To investigate the performance of DICOSAR in real data analysis, we applied DICOSAR to differential co-expression analysis of bulk RNA-seq and snRNA-seq data in the human frontal cortex. We focus on identifying genes that show differential co-expression with *APOE* between control and AD groups because *APOE* is the top risk factor of AD and expresses abundantly in multiple neural cell types. We used the PCC of normalized expression to measure the co-expression between two genes, and therefore detecting differential co-expression amounts to testing the equality of the PCCs between the two groups. In the first analysis, we tested the equality of correlations between the normalized expression of *APOE* and that of each of the 23,535 genes in a bulk RNA-seq data set comprising 482 samples in ROSMAP. The top gene *SERPINA5* had a p-value of 7.9E-05. Fig. 4B shows that more genes had a larger co-expression with *APOE* in the AD group. Genes whose expression are highly correlated with *APOE* generally showed less differential co-expression between the groups (i.e., most light blue points in the plot are concentrated in the middle in Fig. 4B). We found no significant differential co-expressed genes after the multiple testing correction based on either 5% familywise error rate using Bonferroni correction or 5% false discovery rate using the Benjamini-Hochberg procedure (Benjamini and Hochberg, 1995). However, the upward trend in the lower part of the p-value distribution suggests that a large number of genes may be differentially co-expressed with *APOE* between the two groups (Fig. 4A). The lack of significant findings can result from the limited sample size because a large sample size is required for a decent statistical power as shown in our simulation study (Fig. 2). The p-values from DICOSAR are highly consistent (PCC=0.998) with those from the pooled residual permutation performed on the same data set (Fig. 4C).

**Figure 4.**
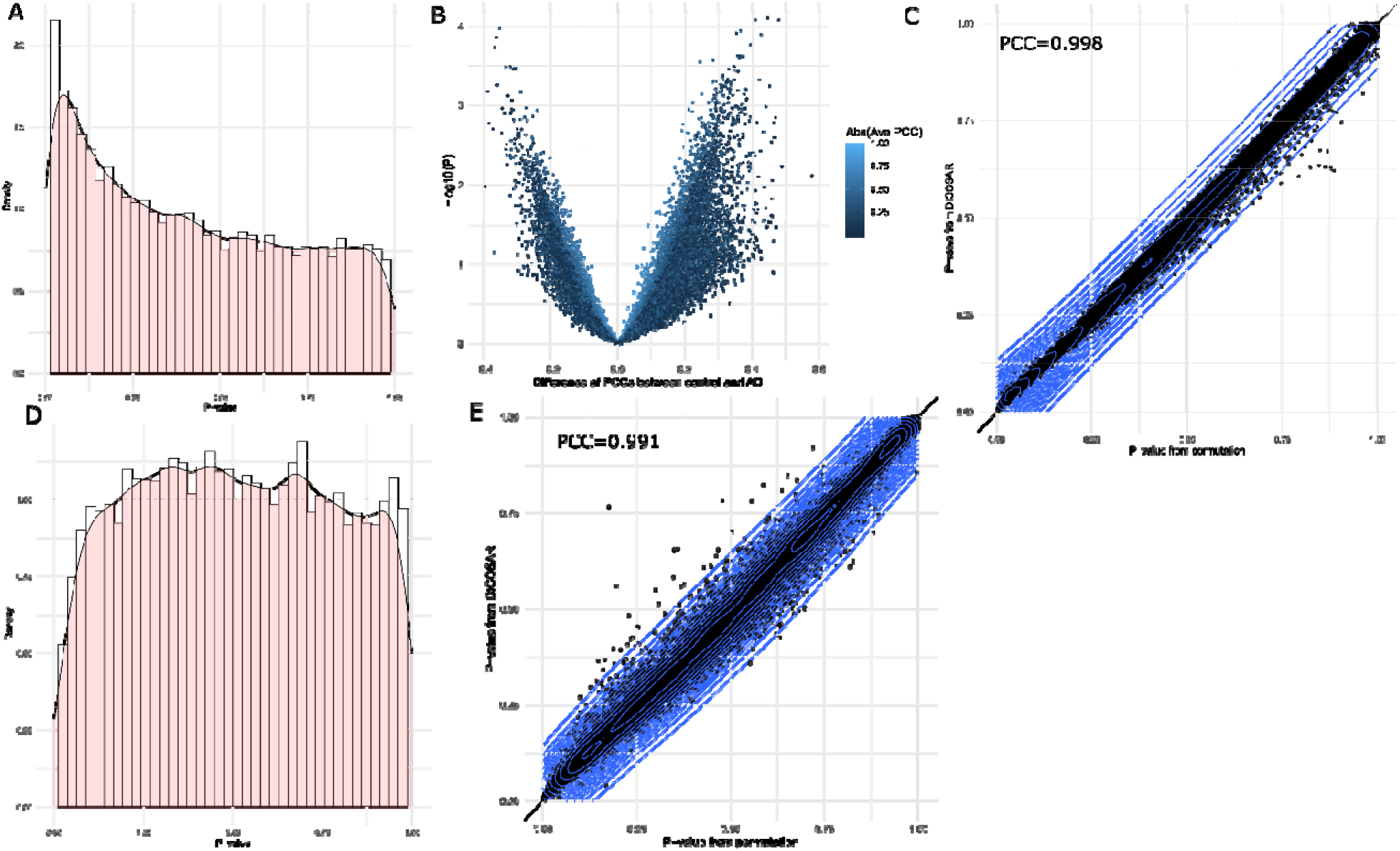
Performance of DICOSAR in the real data differential co-expression analysis. (A)-(C): (A) The p-value distribution, (B) the difference of the PCCs between the AD and control groups (PCC in AD – PCC in control) versus the minus logarithm of the p-values, and (C) a comparison of the p-values estimated by the pooled residual permutation with those estimated by DICOSAR from the differential co-expression analysis of *APOE* with the 482-sample bulk RNA-seq data in ROSMAP. Abs (Ave PCC): the absolute value of the average PCC of the two groups. (D) & (E): (A) The p-value distribution and (B) a comparison of the p-values estimated by the pooled residual permutation with those estimated by DICOSAR from the differential co-expression analysis of *APOE* with the 48-sample snRNA-seq RNA-seq data in ROSMAP.

In the second analysis, we performed a similar differential co-expression analysis for 16,572 genes (after removing very low-expression genes) in astrocytes for *APOE* using a snRNA-seq data set in ROSMAP comprising 48 samples. We observed a flat p-value distribution and a marked dip at the lower end of the distribution of the p-values (Fig. 4D). This is consistent with the deflation of empirical type I error rate observed under the small sample size of 25 in the simulation study, leading to a lack of power to detect very significant differential correlations. The p-values between DICOSAR and the pooled residual permutation are still consistent (PCC=0.991) (Fig. 4E) but to a lesser degree that that seen in the bulk data. This is probably due to the much smaller sample size and the fact that the snRNA-seq data set is sparser than the bulk data set and the distribution of most genes has larger skewness.

### DICOSAR is computationally efficient

After demonstrating its robust statistical performance, we finally evaluated the computational efficiency of DICOSAR. We compared DICOSAR with the pooled residual permutation methods, with 500 and 5000 replicates, respectively. We benchmarked their computational time for testing the equality of the PCCs between two groups of a sample size ranging from 25 to 800. The computational time of DICOSAR for a single test for equal PCCs is comparable to that of the permutation with 500 replicates and is ~10-fold faster than that of permutation with 5000 replicates (Fig. 5). The computational burden of DICOSAR increases at the same rate as the permutation with the increasing sample size.

**Figure 5.**
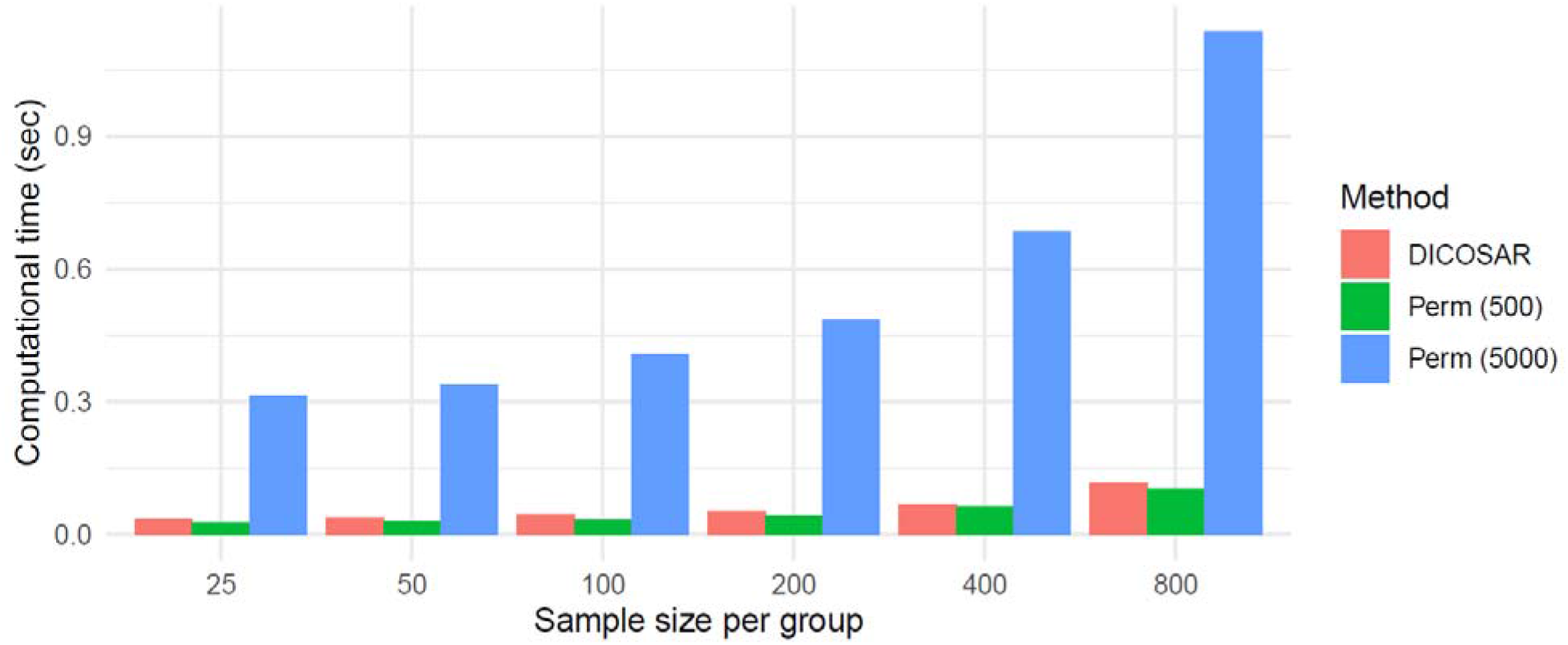
The computational time of DICOSAR for a single test for the equality of the PCCs of a pair of variables between two groups and its comparison with that of the permutation methods. The benchmark is computed based on the average computational time across 20 randomly generated data sets. Perm (500) & Perm (5000): the pooled residual permutation method with 500 and 5000 replicates, respectively.

## Discussion

In this work, we developed DICOSAR, a robust method for a two-sample test for the equality of PCCs. The major advantages of the proposed method include its accuracy under a small sample size, its robustness against the violation of the normality assumption upon the tested variables and its computational efficiency in large-scale studies since it does not depend on a resampling method. Our simulation study demonstrates that DICOSAR is comparable to the permutation and controls the type I error rate substantially better than the Delta method and the separate bootstrap method. This is not surprising that the Delta method performs the worst among these methods because it assumes that both the summary statistics and the test statistic follow normal distributions, and such an assumption does not hold if the sample size is small and either variable is not from a normal distribution. In contrast, the separate bootstrap method only assumes that the test statistic follows a normal distribution, which is less restrictive than the Delta method. Nevertheless, under a small sample size, the z-transformation does not follow a normal distribution generally, and this is the reason for the better performance of DICOSAR than the separate bootstrap. The simulation results also suggest that, compared to the simple approximation (8), the improvement of the higher-order approximation (6) is impressive particularly for small sample sizes.

The co-expression analyses of *APOE* using the bulk RNA-seq data does not identify a significant gene although we did observe a deviation of the p-value distribution from the null distribution. One reason can be that the tests are correlated rather than independent. Because the expression levels of many genes, e.g., cell-type marker genes, are strongly correlated with each other at the bulk tissue level, the actual number of tests are much lower and thus the general Bonferroni correction or the Benjamini-Hochberg procedure (Benjamini and Hochberg, 1995) might be too conservative. Another reason is that this sample size is probably not enough to achieve more significant p-values because of its limited statistical power. As shown in the simulation study, detecting differential correlation requires a large sample size to achieve a decent statistical power. Another approach for detecting differential co-expression is to use a regression model, e.g., NEBULA (He et al., 2021), with an interaction term between the expression and the group. Nevertheless, an interaction model often requires a large sample size as well.

The algorithm of DICOSAR can be expanded in multiple ways to handle more complicated situations. In the current work, we only consider a two-sample test in DICOSAR. An extension to a multi-sample test is possible by modifying the test statistic to aggregate the evidence from more than two groups. For example, the sum of squared differences 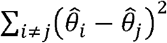 can be such a statistic. In addition, the pooled residual bootstrap is adopted in DICOSAR as the distribution approximated by the multivariate saddlepoint method. As aforementioned, despite being able to better control the type I error rate and borrow information from both groups, this strategy assumes that the two groups share higher moments or at least the joint fourth moments asymptotically. This is because the variance of the estimate of the second moments asymptotically depends on the fourth moments as shown in (Zhang and Boos, 1992, 1993). The simulation results show that a deviation of this assumption can lead to some inflation of type I error rate. This means that a rejection can result from either the heterogeneity of the correlation matrices or higher moments of the distributions between the two groups. If the rejection due to the latter is a major concern, a separate residual bootstrap scheme proposed in (Yang and DeGruttola, 2012; Zhang and Boos, 1993) can be used, which relax the assumption of the shared fourth moments. Extension of DICOSAR to this separate bootstrap scheme is straightforward by applying the saddlepoint approximation to the conditional distribution of the transformed residuals in each group separately.

In summary, DICOSAR is an accurate and robust statistical method for detecting differential correlation and provides a fast alternative to the permutation method. It can also be used for testing differential correlation matrices. DICOSAR provides an analytical approach to facilitate such analyses at a large scale.

## Acknowledgements

This research was supported by Grants from the National Institute on Aging (R01 AG065477, R01 AG070488, and R01 AG061853). The funders had no role in study design, data collection and analysis, decision to publish, or manuscript preparation. The content is solely the responsibility of the authors and does not necessarily represent the official views of the National Institutes of Health.

## Author contributions

I.P. and L.H. conceived the algorithms. L.H. and I.P. developed and implemented the method. L.H. and S.W. performed the simulation study and analyzed the real data. A.K. contributed to acquiring the real data, and discussion of results. All authors contributed to the writing of the manuscript.

## Supplementary figures

**Figure S1:**
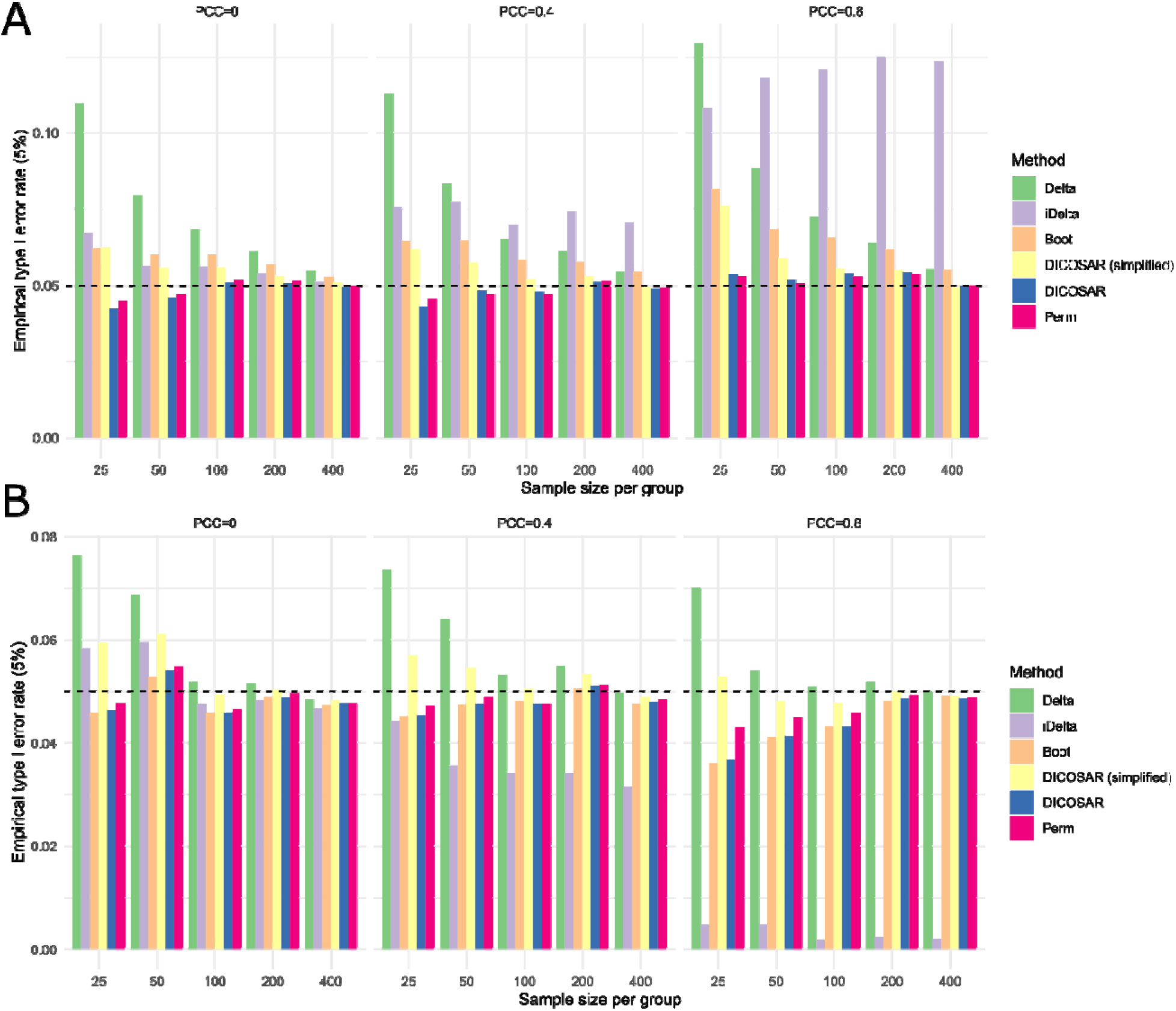
Empirical type I error rate of six methods at the significance level of 5% for testing the equality of PCCs between two groups. The data are generated from a linear transformation of independent variables from (A) a Student’s t-distribution with 6 d.f. and (B) a mixture of two normal distributions. Delta: the Delta method; iDelta: the improved Delta method; Boot: the bootstrap method; Perm: the pooled residual permutation; DICOSAR (simplified): a simplified version of DICOSAR using the approximation (8) instead of (6).

**Figure S2:**
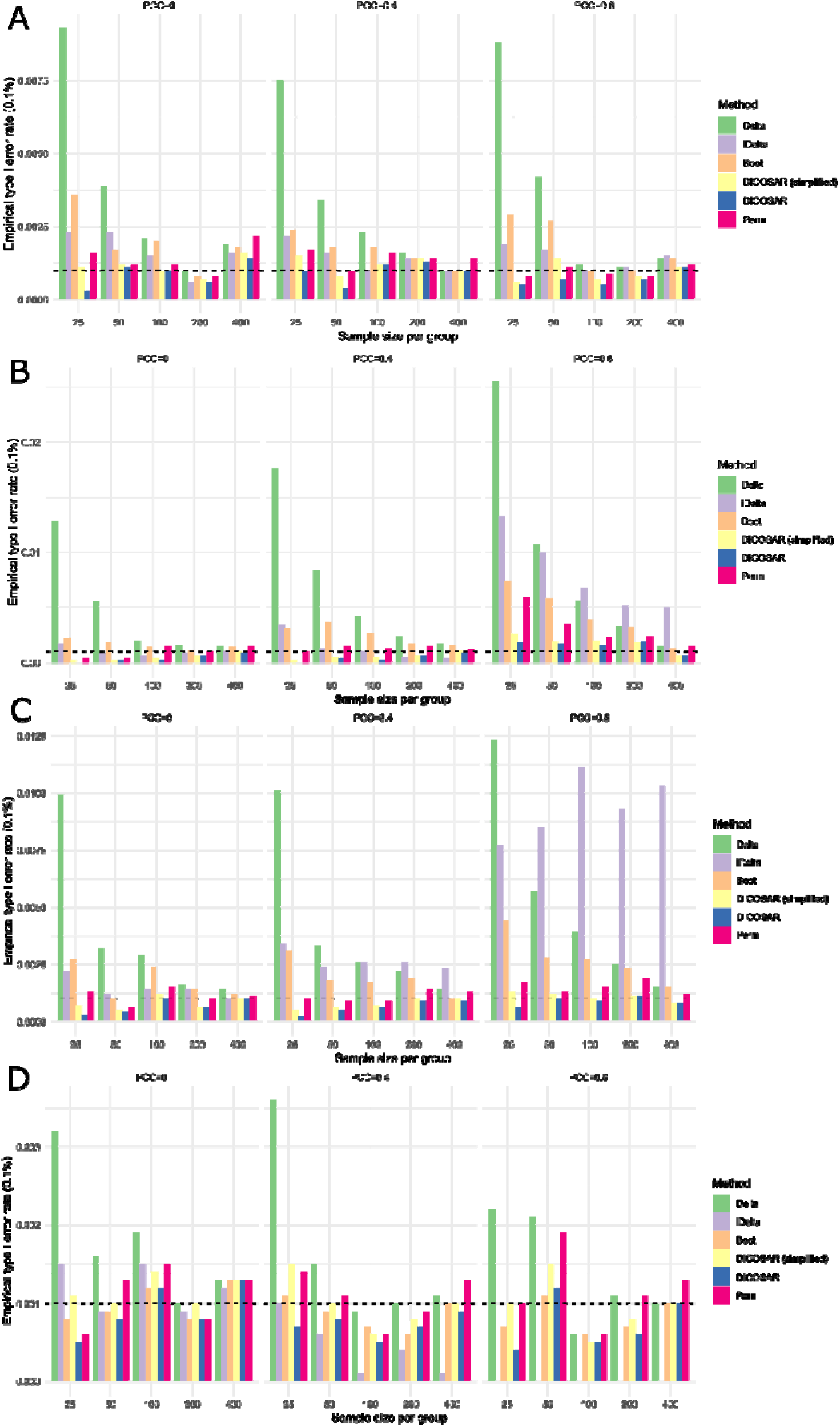
Empirical type I error rate of six methods at the significance level of 0.1% for testing the equality of PCCs between two groups. The data are generated from a linear transformation of independent variables from (A) a standard normal distribution, (B) a gamma distribution with the shape and rate equal to 1, (C) a Student’s t-distribution with 6 d.f. and (D) a mixture of two normal distributions. Delta: the Delta method; iDelta: the improved Delta method; Boot: the bootstrap method; Perm: the pooled residual permutation; DICOSAR (simplified): a simplified version of DICOSAR using the approximation (8) instead of (6).

**Figure S3.**
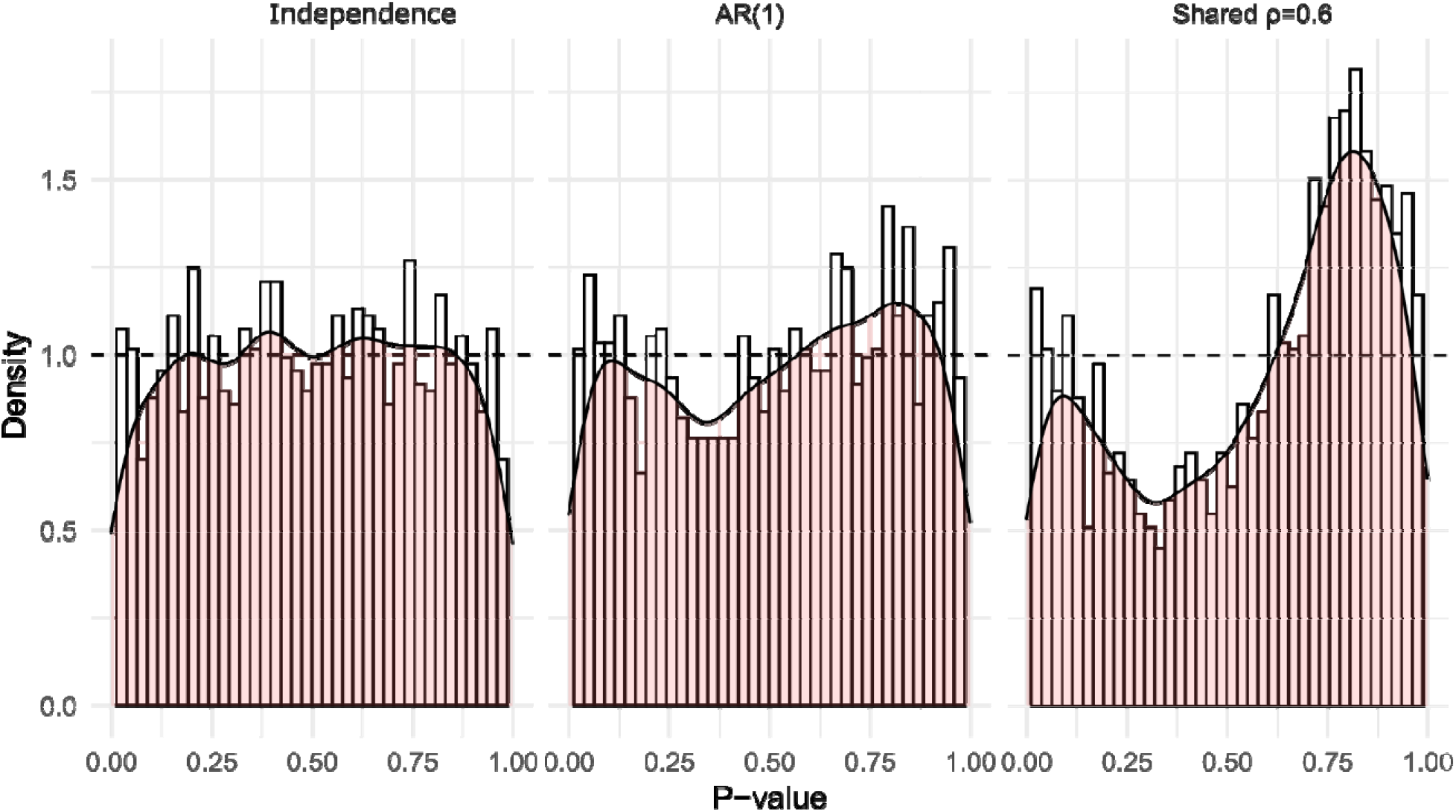
Distribution of p-values from the global test for the equality of two correlation matrices of different structures using the CCT. In each of the structures, 1000 p-values are computed under the null hypothesis. The dimension of the correlation matrices is 10.

